# Cetacean Mammals of the Black and Azov Seas as Indicators of Habitat Quality via Stacked Species Distribution Models

**DOI:** 10.64898/2026.07.07.736995

**Authors:** V. Tytar, L. Fedorenko

**Affiliations:** I.I. Schmalhausen Institute of Zoology, National Academy of Sciences of Ukraine Kyiv, Ukraine; Institute of Hydrobiology, National Academy of Sciences of Ukraine Kyiv, Ukraine

**Keywords:** habitat quality, stacked species distribution models (SSDM), ensemble species distribution models (ESDMs), Black Sea cetaceans, species richness, environmental drivers, conservation prioritisation, marine spatial planning

## Abstract

Habitat degradation and biodiversity loss in the Black and Azov Seas necessitate improved tools for spatially explicit conservation planning. We employed stacked species distribution modelling (SSDM) to assess habitat quality for the three resident cetacean species—the common dolphin (*Delphinus delphis ponticus*), the bottlenose dolphin (*Tursiops truncatus ponticus*), and the harbour porpoise (*Phocoena phocoena relicta*)—which serve as apex predators and indicators of ecosystem health. Occurrence data were compiled from the Global Biodiversity Information Facility (GBIF), and ensemble species distribution models (ESDMs) were constructed using nine algorithms within the SSDM framework, with eight environmental predictors extracted from Bio-ORACLE v3.0. Individual ESDMs demonstrated excellent predictive performance (AUC: 0.82–0.83; TSS: 0.65–0.67; prop.correct: 0.82–0.83). However, the initial continuous stacking method (*pSSDM*) yielded low community-level prediction success (0.36), prompting evaluation of three correction approaches. The *Probability Ranking Rule (PRR)* substantially improved performance (prediction.success = 0.459, sensitivity = 0.704, Jaccard = 0.465), effectively mitigating the overprediction bias inherent in stacked models. Species richness mapping identified multi-species hotspots along the southwestern Black Sea shelf, the Crimean coast, the Kerch Strait, and parts of the eastern coast, while the deep central basin exhibited the lowest richness. Variable importance ranking revealed bathymetry as the primary community-level driver (41.2%), followed by dissolved oxygen (13.8%), sea surface temperature (11.9%), and salinity (10.4%). Species-specific importance patterns confirmed ecological niche segregation, with common dolphins favouring deeper offshore waters and bottlenose dolphins and harbour porpoises associated with shallower shelf environments. The moderate richness observed in the highly productive northwestern shelf, despite high nutrient inputs, may reflect a combination of natural factors (elevated turbidity, reduced salinity) and anthropogenic pressures (fisheries bycatch, shipping, coastal development, and military activity) that limit species co-occurrence. Our findings demonstrate that *PRR*-corrected SSDM provides a robust framework for mapping cetacean habitat quality and identifying conservation priorities in the Black and Azov Seas, offering an evidence-based tool to inform ecosystem-based management in this ecologically unique and increasingly pressured marine region.

## Introduction

### 1. The Imperative for Habitat Quality Assessment

Habitat degradation and biodiversity loss represent two of the most pressing environmental challenges of the Anthropocene, with profound consequences for ecosystem functionality, resilience, and human well-being (Panigada et al., 2024). Despite international commitments—ranging from the Aichi Biodiversity Targets to the EU Biodiversity Strategy to 2020—conservation targets remain largely unmet at both global and regional scales. This persistent shortfall underscores the need for improved, cost-effective tools for biodiversity monitoring and habitat quality assessment that can bridge the gap between science and policy.

Habitat quality (HQ)—the capacity of an ecosystem to provide suitable conditions for the persistence of species—is a critical metric for conservation planning. Spatially explicit assessments of HQ enable the identification of priority areas for protection, the evaluation of anthropogenic impacts, and the optimization of conservation interventions (Wang et al., 2024; Yao et al., 2025). Among the various approaches to HQ mapping, species distribution models (SDMs) and expert-based models have emerged as complementary tools, with SDMs generally demonstrating superior predictive accuracy when high-quality occurrence data are available (Taufiq, Nurainas, 2025).

The Black and Azov Seas present unique challenges for habitat quality assessment due to their semi-enclosed nature, low salinity gradients (Azov: 9-14‰), and history of eutrophication and biological invasions (Zaitsev, Mamaev, 1997; Zaitsev et al., 2002; ANEMONE, 2021). These unique oceanographic conditions create a strongly stratified water column with a permanent halocline and a large anoxic zone, which profoundly influence the distribution of marine life and ecosystem functioning. The three cetacean species resident in the Black Sea—the common dolphin (*Delphinus delphis ponticus*), the bottlenose dolphin (*Tursiops truncatus ponticus*), and the harbour porpoise (*Phocoena phocoena relicta*)—are apex predators and therefore serve as important indicators of overall ecosystem health. Previous research has established that their distributions are strongly linked to key environmental gradients, with common dolphins preferring deeper offshore waters, while harbour porpoises and bottlenose dolphins are more frequently associated with shallower shelf areas. Significant relationships have been identified between their occurrence and parameters such as sea surface temperature and bathymetry, suggesting a degree of niche segregation. However, comprehensive basin-wide assessments of their habitat suitability that explicitly rank the environmental drivers of their distribution remain scarce.

To address this gap, the present study employs the Stacked Species Distribution Modelling (SSDM) framework (Schmitt et al., 2017), which offers a robust and flexible approach to habitat quality assessment through the construction of ensemble species distribution models (ESDMs). The ensemble approach is particularly advantageous because it integrates predictions from multiple algorithms, rather than relying on a single model. This multi-algorithm strategy mitigates the limitations and biases inherent in any individual method, substantially reducing prediction uncertainty and enhancing overall model accuracy and transferability. By combining the strengths of diverse modelling techniques, ensemble models consistently outperform single-algorithm models, providing more reliable habitat suitability maps that are essential for evidence-based conservation decision-making. Furthermore, the SSDM framework facilitates the stacking of individual species predictions to generate maps of species richness and community composition, while simultaneously enabling the quantification and ranking of predictor variable importance—a critical feature for identifying the primary environmental drivers shaping cetacean distributions in this ecologically complex marine system.

The aim of this study is to conduct a spatially explicit assessment of habitat quality for the three Black and Azov sea cetacean species and to identify and rank the most influential environmental factors driving their distributions. This will be achieved by constructing robust ensemble species distribution models within the SSDM framework using global environmental data from the Bio-ORACLE v3.0 dataset.

To achieve this aim, we have set the following tasks:

1. ***Data Compilation and Preparation***. Compile and standardize occurrence records for the three cetacean species (*D. delphis ponticus, T. truncatus ponticus*, and *P. phocoena relicta*) from available databases and literature. Concurrently, extract a comprehensive set of environmental predictor variables from the Bio-ORACLE v3.0 dataset for the entire Black and Azov Seas study area. All data will be gridded to a common spatial resolution to ensure compatibility for modelling.
2. ***Ensemble Model Construction***. For each of the three cetacean species, build ensemble species distribution models (ESDMs) by running multiple algorithms within the SSDM framework. This diversity of algorithms is expected to ensure that both parametric and non-parametric relationships between species occurrences and environmental predictors are adequately captured. Model calibration will be conducted using cross-validation to ensure robustness. The ensemble prediction for each species will be generated as a weighted average of individual algorithm outputs, with weights determined by model performance metrics (Allouche et al., 2006; Boyce et al., 2002; and others), ensuring that better-performing models contribute more to the final ensemble prediction.
3. ***Habitat Quality Mapping and Species Richness Stacking***. Generate high-resolution habitat suitability maps for each of the three species across the entire study area. Subsequently, apply the stacking module of the SSDM framework to sum the individual species occurrence probabilities, producing a map of predicted species richness for the Black and Azov Seas. This provides a spatially explicit assessment of biodiversity hotspots and habitat quality at the community level.
4. ***Environmental Driver Ranking***. Utilize the variable importance functions within the SSDM framework to quantify and rank the relative contribution of each environmental predictor variable to the ensemble models for each species. This will reveal the primary environmental gradients driving the distribution of each cetacean species, distinguishing between key factors.
5. ***Model Evaluation and Ecological Interpretation***. Critically evaluate the predictive performance of all individual algorithms and the final ensemble models using independent evaluation metrics. Interpret the resulting habitat suitability and species richness maps in the context of known ecological preferences, identified niche segregation patterns, and contemporary conservation challenges facing Black Sea cetaceans, thereby translating model outputs into actionable conservation insights.

By employing this rigorous ensemble-based SSDM methodology, this study aims to deliver a robust, transparent, and ecologically meaningful assessment of habitat quality that can directly inform conservation planning and policy development for these vulnerable marine mammal populations.

Finally, marine habitat mapping, particularly when integrated with advanced species distribution modelling, is essential for implementing ecosystem-based management in the Black and Azov Seas. The existing data on cetaceans provides a strong foundation for our proposed work. By building a stacked SDM, one can create a critical spatial tool that visualizes the distribution of all three cetacean species simultaneously. This will help identify areas of high conservation value, predict potential overlaps with human activities (like shipping lanes or fishing grounds), and ultimately support the prioritization of areas for protection. This approach is exactly the kind of evidence-based, spatially explicit planning that is urgently needed to safeguard these unique and pressured marine ecosystems.

## 2. Methods

### 2.1. Occurrence data

The use of stacked SDM is well-aligned with recent, basin-wide research. The most comprehensive data comes from the EU-funded CeNoBS project, which conducted a large-scale aerial survey across the Black Sea in 2019, covering over 7,000 km of transects (Paiu et al., 2024). This landmark survey yielded 1,744 cetacean sightings and provided the first robust basin-wide abundance estimates for all three Black Sea subspecies. However, while the CeNoBS dataset offers unparalleled spatial coverage for a single time period, it is constrained to summer months and represents a single-year snapshot.

To capture a more comprehensive picture of species distributions across seasonal and interannual variability, the present study ultimately draws occurrence data from the Global Biodiversity Information Facility (GBIF). GBIF serves as the world’s largest and most comprehensive biodiversity database, aggregating occurrence records from hundreds of datasets worldwide. For the Black Sea region, GBIF compiles multiple historical and ongoing cetacean monitoring efforts, including dedicated aerial and vessel-based surveys, stranding records, and systematic monitoring programmes. The key GBIF datasets relevant to this study include:

- The “Cetacean sightings in the Black Sea, Sea of Azov and Kerch Strait” dataset, containing over 645 sightings collected between 1993 and 2010 through dedicated line-transect aerial and vessel-based surveys;
- The “Cetacean strandings in the northern Black Sea and the Sea of Azov” dataset, recording more than 1,220 stranding events along the Crimean coast from 1989 to 2010;
- The “Green Balkans NGO’s cetacean sightings in the Bulgarian Black Sea waters”, comprising 1,663 occurrences from vessel surveys conducted between 2015 and 2022.

By aggregating these diverse sources, GBIF provides a temporally and spatially extensive occurrence dataset that captures the full known distribution of the three Black & Azov sea cetacean subspecies (Table 1), thereby maximising the robustness and generalisability of the ensemble SDMs.

**Table 1.** Occurrences representing the distribution of Black & Azov sea cetacean subspecies.

| Subspecies | Number of Occurrences | GBIF Occurrence Download |
| --- | --- | --- |
| *D. delphis ponticus* | 1,223 | https://doi.org/10.15468/dl.t9uecn |
| *T. truncatus ponticus* | 1,470 | https://doi.org/10.15468/dl.6k53b9 |
| *P. phocoena relicta* | 5,507 | https://doi.org/10.15468/dl.97dwv7 |

### 2.2. Environmental data

To characterise the environmental niche of the three Black Sea cetacean species and to construct robust ensemble species distribution models, we selected a suite of eight ecologically relevant predictor variables from the Bio-ORACLE v3.0 database (Assis et al., 2024). Bio-ORACLE v3.0 is a globally comprehensive, high-resolution dataset of marine environmental layers specifically designed for species distribution modelling and climate change impact assessments. It provides global coverage at a spatial resolution of 0.05° (approximately 5.5 km) and includes both surface and benthic layers derived from satellite observations and oceanographic models. For the present study, we extracted present-day baseline layers corresponding to the period 2000–2014, which provide a representative contemporary climatology for the Black and Azov seas.

The selection of predictor variables was guided by known ecological preferences of the target species, the unique oceanographic characteristics of the study region, and previous modelling efforts on cetacean distributions in the Black Sea (Paiu et al., 2024; Panigada et al., 2024). The chosen variables encompass key physical, chemical, and biological dimensions of the marine environment that are hypothesised to influence cetacean habitat suitability:

1. *Bathymetry* — measured in metres (m). Bathymetry is a primary driver of cetacean distribution, reflecting the distinct habitat preferences observed among the three species: common dolphins typically occupy deeper offshore waters, while bottlenose dolphins and harbour porpoises are more frequently associated with shallower shelf areas (Panigada et al., 2024; Paiu et al., 2024). This variable also serves as a proxy for distance to coast and seabed topography, which influence prey aggregation.
2. *Sea Surface Temperature* — measured in degrees Celsius (°C). Temperature is a fundamental determinant of species distribution, influencing metabolic rates, reproductive cycles, and the distribution of prey species. Black Sea cetaceans exhibit seasonal movements linked to thermal gradients, and sea surface temperature has been identified as a significant predictor of their occurrence (Panigada et al., 2024).
3. *Salinity* — measured in Practical Salinity Units (PSS). Salinity is particularly relevant for the Black and Azov Seas due to their semi-enclosed nature and pronounced brackish gradients (Azov Sea: 9– 14‰) (Zaitsev & Mamaev, 1997). Salinity influences prey community composition and may act as a direct physiological constraint or an indirect driver through its effects on fish distribution.
4. *Dissolved Molecular Oxygen* — measured in moles per cubic metre (mol/m^3^). This variable is critically important in the Black Sea, where a permanent halocline creates a stratified water column with an extensive anoxic zone below approximately 150–200 m (Zaitsev et al., 2002). Cetaceans are obligate air-breathers confined to oxygenated surface and intermediate waters, making dissolved oxygen a potentially strong predictor of habitat suitability.
5. *Net Primary Productivity* — measured in grams of carbon per cubic metre per day (g/m^3^/day). Primary productivity serves as a proxy for food web productivity; higher productivity areas support greater fish biomass, which in turn attracts foraging cetaceans. This variable is particularly relevant for understanding seasonal and spatial variation in habitat quality.
6. *Chlorophyll-a Concentration* — measured in milligrams per cubic metre (mg/m^3^). Chlorophyll-a is a direct indicator of phytoplankton biomass and, by extension, potential foraging hotspots. It has been widely used in cetacean habitat modelling as a proxy for trophic resource availability (Paiu et al., 2024).
7. *Mixed Layer Depth* — measured in metres (m). The mixed layer depth influences the vertical distribution of prey species and, consequently, foraging efficiency of cetaceans. A shallow mixed layer may concentrate prey near the surface, while deeper mixing can disperse prey throughout the water column, affecting encounter rates and habitat use.
8. *Sea Water Velocity* — measured in metres per second (m/s). Currents play a significant role in prey aggregation through frontal systems, upwelling zones, and eddy formation. Cetaceans are often observed foraging along current boundaries where physical processes concentrate plankton and fish, making this variable a valuable predictor of suitable foraging habitat.

All environmental layers were extracted for the entire Black and Azov seas study area using the ‘sdmpredictors’ R package, which provides seamless access to Bio-ORACLE v3.0 data (Bosch et al., 2023). To avoid multicollinearity issues that can compromise model performance—particularly for algorithms such as Generalised Linear Models—we conducted a correlation analysis among the eight predictor variables using Pearson’s correlation coefficient. Variables exhibiting a correlation coefficient of |r| > 0.7 were considered for exclusion or combination, following established protocols in SDM studies (Dormann et al., 2013). The final predictor set was retained after ensuring that the variance inflation factor (VIF) for each variable remained below a threshold of 10, indicating acceptable levels of multicollinearity (Montgomery et al., 2021). Under these conditions *Chlorophyll-a Concentration* was excluded as far as the VIF for this variable considerably exceeded the adopted threshold and in addition strongly correlates with *Net Primary Productivity* (r>0.9).

All variables were projected using SAGA GIS (Conrad et al., 2015) to the study area extent for subsequent modelling.

### 2.3. Model Calibration and Ensemble Construction

Species distribution models were constructed for each of the three cetacean species using the SSDM package in R (Schmitt et al., 2017), which provides a comprehensive framework for integrating multiple algorithms and generating consensus predictions. The modelling workflow proceeded through three hierarchical levels: (1) individual Species Distribution Models (SDMs) fitting occurrences of a single species using a single algorithm; (2) Ensemble SDMs (ESDMs) combining the outputs of several SDMs, each employing a different modelling algorithm; and (3) the Stacked SDM (SSDM) combining ESDM outputs to model species assemblages and compute species richness. Algorithms used include Generalized linear model (*GLM*), Generalized additive model (*GAM*), Multivariate adaptive regression splines (*MARS*), Generalized boosted regressions model (*GBM*), Classification tree analysis (*CTA*), Random forest (*RF*), Maximum entropy (*MAXENT*), Artificial neural network (*ANN*), and Support vector machines (*SVM*).

Following the calibration of individual SDMs, we constructed Ensemble Species Distribution Models (ESDMs) for each of the three species by combining the predictions of all four algorithms. The ensemble approach is methodologically superior to single-algorithm modelling because it mitigates the limitations and biases inherent in any individual method, substantially reducing prediction uncertainty and enhancing overall model accuracy and transferability. Two consensus methods are implemented in the SSDM package for combining SDM outputs: a simple average and a weighted average. We adopted the weighted average consensus, where weights are assigned to each algorithm based on a user-specified metric—in this case, a combined performance score derived from AUC and TSS values. This ensures that better-performing models contribute more to the final ensemble prediction.

### 2.4. Variable Importance Ranking

A central objective of this study is to identify and rank the environmental factors most influential in shaping the habitat suitability of each cetacean species. The SSDM framework provides built-in functionality for quantifying the relative contribution of each predictor variable to the ensemble models. Variable importance is calculated for each individual SDM algorithm, and the resulting importance scores are then aggregated to provide a robust, multi-algorithm consensus ranking.

The final variable importance ranking for each species is generated by averaging the importance scores across all individual algorithms, weighted by each algorithm’s performance metric (AUC or TSS). This produces a consensus ranking that identifies the primary environmental drivers of cetacean distribution in the Black and Azov Seas.

### 2.5. Species Richness Stacking and Habitat Quality Mapping

Finally, after constructing ESDMs for each of the three species, we applied the stacking module of the SSDM framework to generate a basin-wide assessment of species richness. We employed the continuous habitat suitability map stacking method (pSSDM), which sums the continuous occurrence probabilities of individual species to produce a map of predicted local species richness. However, there is key challenge in community-level modeling: excellent single-species models can still produce second-rate predictions when stacked. The most common and well-documented cause for this is overprediction of species richness (Calabrese et al., 2014). In this respect the SSDM package offers some built-in options that can be useful.

1. *bSSDM* (*Binary Stacked SDM*). Instead of summing probabilities, this method first converts each species’ continuous habitat suitability map into a binary presence/absence map based on a threshold. It then sums these binary maps.
2. *MaximumLikelihood*. This method uses a statistical approach to adjust the species richness predicted by the pSSDM to better match the observed richness, effectively “correcting” the overprediction bias.
3. *PRR.pSSDM (Probability Ranking Rule)*. This method is a correction approach. It first predicts species richness (using pSSDM, for example) and then adjusts the individual species predictions to match that richness, effectively ranking species by their habitat suitability and selecting the top ones. This directly addresses the overprediction of composition.

Further we can compare the results from these different methods and see which one yields the highest prediction success and Jaccard index (a measure of community similarity).

This approach allows to conduct a spatially explicit assessment of biodiversity hotspots and habitat quality at the community level, identifying areas of high conservation priority where all three cetacean species are likely to co-occur.

The one notable limitation is the lack of consideration of biotic interactions (like competition) in the standard SSDM workflow, which typically applies only an “environmental filter” (D’Amen et al., 2018). In a community of three top predators, which may have overlapping diets and spatial niches, this is an important point to acknowledge. However, this is a general limitation of the approach and not a reason to abandon its application.

## 3. Results

### 3.1 Model Performance and Evaluation

The ensemble species distribution models demonstrated excellent predictive performance for all three species. AUC values ranged from 0.82 to 0.83, indicating strong discriminatory power. TSS values (0.65–0.67) were well above the 0.4 threshold considered indicative of reliable model performance (Allouche et al., 2006). The proportion of correctly predicted observations was consistently high (82– 83%), confirming that the the multi-algorithm strategy effectively captured the ecological niches of the three considered cetacean subspecies (Table 2).

**Table 2.** Predictive performance metrics of ensemble species distribution models (ESDMs) for the three Black & Azov sea cetacean species.

| Species | AUC* | TSS** | prop.correct*** |
| --- | --- | --- | --- |
| *D. delphis ponticus* | 0.82 | 0.65 | 0.82 |
| *T. truncatus ponticus* | 0.83 | 0.67 | 0.83 |
| *P. phocoena relicta* | 0.83 | 0.65 | 0.83 |
| Interpretation of metrics | Excellent discrimination: the models can distinguish between presences and absences 82–83% of the time across all possible thresholds | Good to very good: the model's ability to correctly classify presences and absences is substantially better than random (TSS > 0.4 is reliable; > 0.6 is good) | Excellent classification accuracy: the overall classification accuracy is very high |
| *Area Under the Curve, **True Skill Statistic, ***proportion of correctly predicted observations |  |  |  |

### 3.2. Stacked Species Distribution Model Performance and Comparison of Stacking Methods

Following the successful construction of individual ensemble species distribution models (ESDMs) for each of the three cetacean species, we applied the stacking module of the SSDM framework to generate community-level predictions of species richness and composition. However, the initial application of the continuous stacking method (*pSSDM*), which sums the continuous occurrence probabilities of individual species, yielded a relatively low prediction.success of 0.36. This result is consistent with a well-documented challenge in stacked species distribution modelling: excellent single-species models can produce suboptimal predictions when stacked due to the accumulation of commission errors, leading to systematic overprediction of species richness (Calabrese et al., 2014).

To address this overprediction bias, we evaluated three alternative stacking methods implemented in the SSDM package: *bSSDM, MaximumLikelihood*, and *PRR.pSSDM* (see above). Each method employs a different approach to correct for the tendency of *pSSDM* to overpredict species occurrences. The *PRR* approach substantially improved community-level performance metrics compared to *pSSDM*. The prediction.success increased from 0.36 to 0.459, while sensitivity (the proportion of true presences correctly identified) reached 0.704, indicating that the model successfully captures more than 70% of true species occurrences. The Jaccard index of 0.465 reflects moderate similarity between predicted and observed community composition. The species richness error of 0.572 indicates that, on average, the model is off by approximately 0.57 species per site—representing about 19% of the total species pool (three species)—which is a moderate error. Specificity (0.566) remained moderate, suggesting some residual overprediction of absences, a known limitation of the SSDM approach.

The high sensitivity value (0.704) is particularly noteworthy for conservation applications, as it indicates that the model is effective at detecting actual species occurrences. Failing to identify true presences (false negatives) is generally more critical for conservation planning than overpredicting absences (false positives), as the former may lead to the omission of important habitat areas from protection schemes. Conversely, the moderate specificity (0.566) indicates that approximately 43% of predicted presences are false positives, a residual overprediction that must be considered when interpreting the species richness maps.

The *bSSDM* and *MaximumLikelihood* methods yielded prediction success of 0.355 and 0.354, with the *PRR* approach demonstrating the most favourable balance between sensitivity and specificity. Based on these comparative results, we selected the PRR-corrected SSDM for subsequent habitat quality mapping and variable importance analysis.

### 3.3. Species Richness and Habitat Quality Mapping

The *PRR*-corrected stacked species distribution model generated a spatially explicit map of predicted species richness for the Black and Azov Seas (Figure 1), with colours ranging from red (high values) through orange and yellow to green and blue (low values). This map serves as a community-level indicator of habitat quality, with areas of higher predicted richness (red to orange hues) representing locations where multiple cetacean species are likely to co-occur and where habitat conditions are most favourable for the cetacean community as a whole.

**Figure 1.**
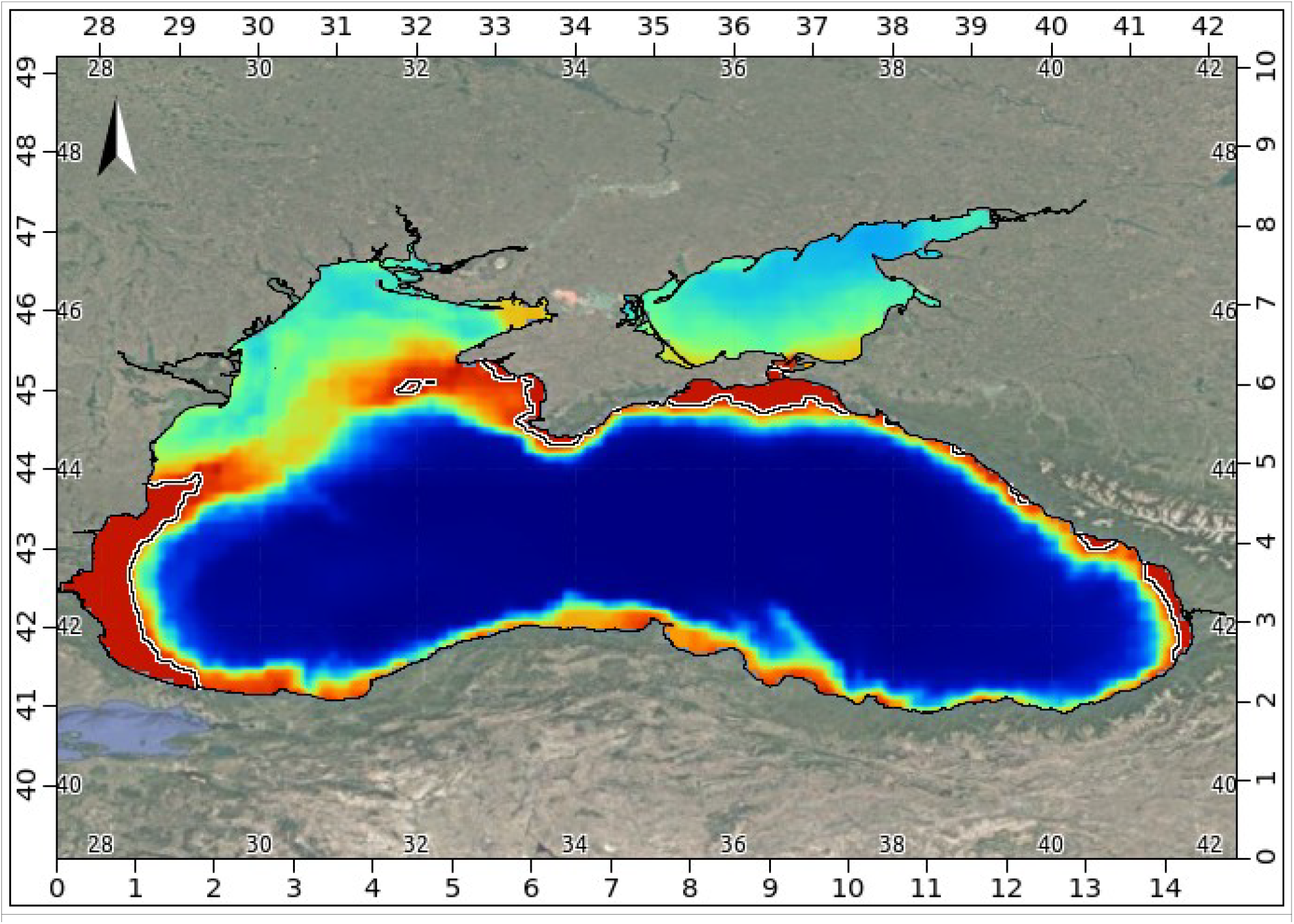
Predicted species richness of cetaceans in the Black and Azov Seas based on the *PRR*-corrected stacked species distribution model. Colours range from red (highest richness, 3 species) through orange, yellow, and green to blue (lowest richness, 0 species). The map depicts a gradient from high richness in productive shelf areas (particularly stretching from the vicinity of the Bosphorus Strait and extending northward along the Turkish and Bulgarian coast), the Crimean coast and Kerch Strait, and parts of the eastern coast) to low richness in the deep central basin. High-richness areas (red to orange) represent multi-species hotspots and priority regions for conservation action. The spatial resolution is 0.05° (approximately 5.5 km). The black&white contour line indicates the 2-species threshold, distinguishing areas with the potential to host multiple cetacean species (core habitat) from those with lower predicted richness.

The species richness map reveals several distinct spatial patterns across the study area.

- *High-richness areas (red to orange hues; 2–3 species)*. The most pronounced high-richness areas are concentrated along the southwestern Black Sea shelf (Bulgarian and Romanian waters) and the coastal waters of the Crimean Peninsula. These regions are characterised by relatively shallow depths (< 200 m), elevated primary productivity, and favourable thermal conditions, supporting the co-occurrence of all three species. The Kerch Strait region, connecting the Black and Azov Seas, also emerges as a high-richness area, possibly reflecting the transitional nature of this hydrographically complex zone. Additional high-richness patches are evident along the eastern coast near the Caucasus, though these areas (particularly to the north of Georgia) are generally more fragmented.
- *Medium-richness areas (yellow to green hues; 1–2 species)*. Extensive areas of the Black Sea, including the northwestern shelf, the western coastal zone, and portions of the eastern shelf, show intermediate species richness (yellow to green). These areas typically support one or two species, most commonly harbour porpoises and bottlenose dolphins in shallower waters or common dolphins in deeper offshore areas. Interestingly, the northwestern Black Sea shelf—a region of high primary productivity supported by nutrient-rich discharges from the Danube, Dnister, and Dnipro rivers—did not emerge as an area of high predicted species richness. Instead, only moderate richness values were observed in this region. This pattern may reflect environmental conditions that are suboptimal for one or more species, such as elevated turbidity from riverine sediment inputs, reduced salinity, or localized anthropogenic pressures. For example, shipping, fishing (Birkun et al.,2014), and coastal development, military activity), all of which may limit the co-occurrence of all three cetacean species in this otherwise productive area. Interestingly, the northwestern Black Sea shelf—a region of high primary productivity supported by nutrient-rich discharges from the Danube, Dnister, and Dnipro rivers—did not emerge as an area of high predicted species richness. Instead, only moderate richness values were observed in this region. This pattern may reflect natural environmental conditions that are suboptimal for one or more species, such as elevated turbidity from riverine sediment inputs or reduced salinity. In addition, the area is subject to numerous anthropogenic pressures, including military activity associated with the ongoing Ukrainian-Russian war, that may further limit the co-occurrence of all three cetacean species in this otherwise productive area (e.g., Birkun et al., 2014; Frassà et al., 2023; Hadzhun, 2023; etc.). The Azov Sea exhibits predominantly medium richness, with harbour porpoises and bottlenose dolphins occurring in this shallow, brackish basin, while common dolphins are largely absent due to the low salinity and shallow depths.
- *Low-richness areas (blue to dark blue hues; 0–1 species)*. The deep central basin of the Black Sea, characterised by depths exceeding 1,000 m, low primary productivity, and the presence of anoxic waters below 150–200 m, shows the lowest predicted species richness (dark blue). This extensive central region forms a prominent blue core extending across the middle of the Black Sea. In these areas, only the offshore-adapted common dolphin is likely to occur, while the shelf-associated bottlenose dolphin and harbour porpoise are largely absent. The low richness in these regions reflects the limited habitat availability for species other than the common dolphin.

#### 3.3.1. Ecological Interpretation

The spatial pattern of high richness concentrated in the southern Black Sea, rather than the northwestern shelf, is noteworthy. While the northwestern shelf is known for high primary productivity due to nutrient inputs from major rivers (Zaitsev et al., 2006), the lower predicted species richness in this region may reflect environmental conditions that are less favourable for the co-occurrence of all three species. Potential factors include the following.

- Reduced salinity — the northwestern shelf receives substantial freshwater inputs, creating a lower-salinity environment that may be less suitable for common dolphins, which prefer more oceanic conditions.
- Elevated turbidity — high suspended sediment loads from rivers may reduce foraging efficiency for visual predators.
- Localised anthropogenic pressures — heavy shipping traffic, fishing activities, and coastal development in this region may reduce habitat quality.
- Specific prey distribution — the prey community structure in the northwestern shelf may support certain species but not others.

The concentration of high richness (red to orange hues) along the southern Black Sea coast (near the Bosphorus and Bulgarian coast) and in the Kerch Strait area suggests that these regions provide the environmental conditions necessary for the co-occurrence of all three species, potentially reflecting a balance of factors that is optimal for the cetacean community. These high-richness areas represent multi-species hotspots where habitat protection would benefit multiple species simultaneously, maximising conservation outcomes. Notably, these areas are subject to significant anthropogenic pressures, including shipping (particularly near the Bosphorus), fishing, and coastal development, underscoring the urgency of implementing protective measures.

### 3.4. Variable Importance Ranking for the Cetacean Community

A central objective of this study was to identify and rank the environmental factors most influential in shaping habitat suitability for the cetacean community as a whole. The SSDM framework provides built-in functionality for quantifying the relative contribution of each predictor variable to the ensemble models. By averaging the importance scores across all individual algorithms and species, weighted by each algorithm’s performance metric, we generated a consensus ranking of environmental drivers for the entire cetacean assemblage (Table 3, Figure 2).

**Table 3.** Relative importance of environmental variables for the Black & Azov Sea cetacean community (averaged across all three species)

| Rank | Environmental Variable | Mean Importance (%) | Interpretation |
| --- | --- | --- | --- |
| 1 | *Bathymetry* | 41.2 | Primary habitat differentiator; distinguishes offshore vs. shelf-associated species |
| 2 | *Dissolved Oxygen* | 13.8 | Critical due to Black Sea anoxia; limits vertical habitat |
| 3 | *Sea Surface Temperature* | 11.9 | Thermal niche; influences prey distribution and metabolic rates |
| 4 | *Salinity* | 10.4 | Brackish gradient influence; particularly relevant for Azov Sea |
| 5 | *Net Primary Productivity* | 8.6 | Proxy for food availability; foraging hotspot indicator |
| 6 | *Mixed Layer Depth* | 8.0 | Affects prey vertical distribution and foraging efficiency |
| 7 | *Sea Water Velocity* | 6.1 | Currents aggregate prey at frontal zones and eddies |

**Figure 2.**
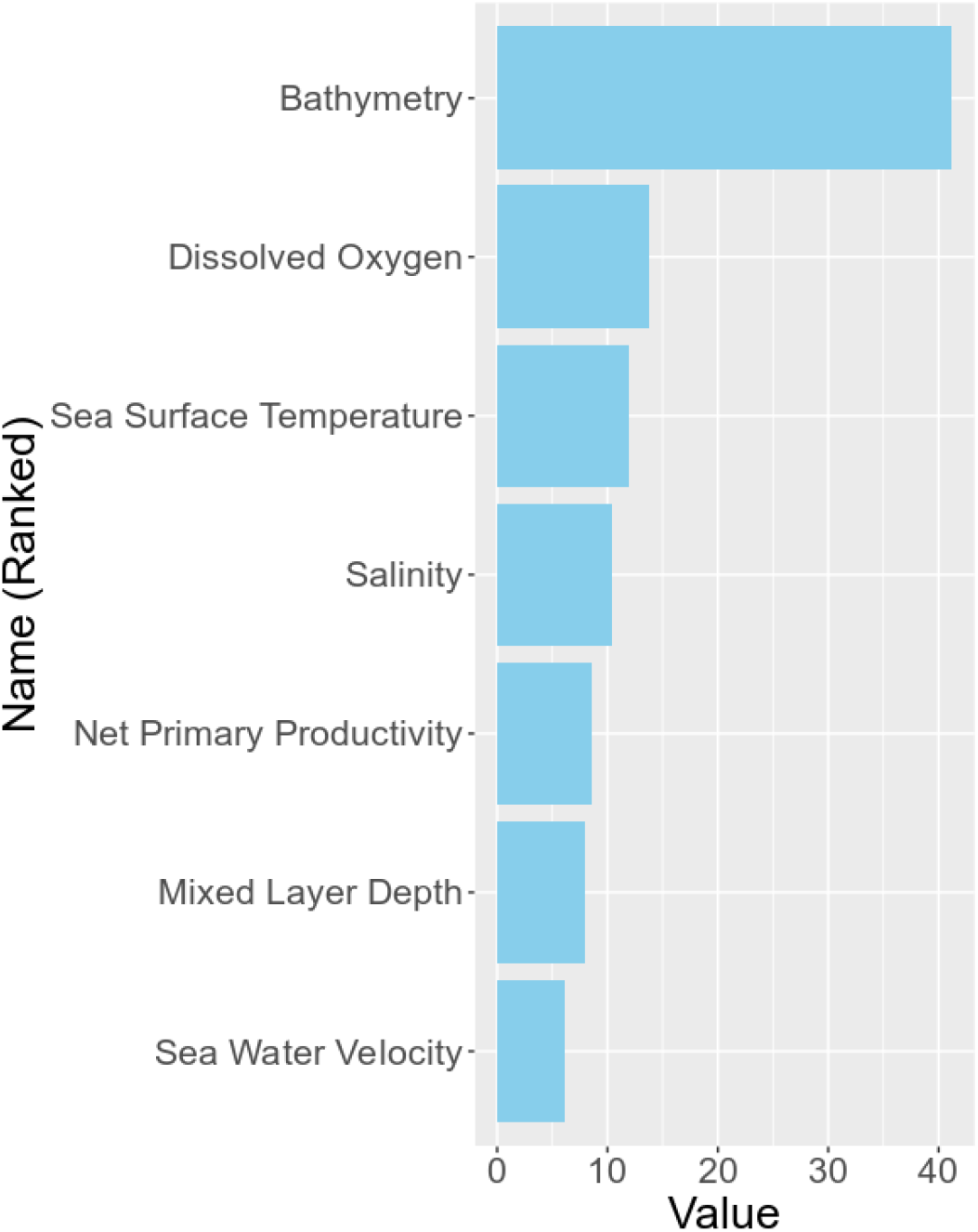
Barplot of relative importance of environmental variables for the Black Sea cetacean community (averaged across all three species)

*Bathymetry* emerged as the most important predictor of cetacean habitat suitability across the community, confirming the well-documented habitat segregation between offshore and shelf-associated species (Panigada et al., 2024; Paiu et al., 2024). This variable serves as a proxy for multiple ecological factors, including distance from coast, seabed topography, and the distribution of prey species. The dominance of bathymetry as a predictor reflects the distinct depth preferences of the three species: common dolphins occupying deeper offshore waters, while bottlenose dolphins and harbour porpoises are associated with shallower shelf areas.

*Dissolved Oxygen* emerged in the second place as a particularly important predictor, highlighting the role of the Black Sea’s unique oxygen stratification in limiting habitat availability. The presence of a permanent anoxic zone below approximately 150–200 m restricts cetacean habitat to oxygenated surface and intermediate waters (Zaitsev et al., 2002). This variable is especially relevant for the harbour porpoise, which tends to occupy shallower waters but may be indirectly affected by oxygen availability through its influence on prey distribution.

*Sea Surface Temperature* ranked as the third most important community-level driver, reflecting the thermal preferences of the species and their seasonal movements in response to thermal gradients.

Temperature influences metabolic rates, reproductive cycles, and the distribution of prey species, making it a fundamental determinant of cetacean habitat suitability.

*Salinity* in the fourth place showed moderate importance, reflecting the influence of brackish conditions, particularly in the Azov Sea (9–14‰) and near river mouths. While cetaceans are generally euryhaline, salinity influences prey community composition and may act as an indirect driver through its effects on fish distribution.

*Net Primary Productivity* ranked next as a proxy for food web productivity, with higher productivity areas supporting greater fish biomass and consequently attracting foraging cetaceans. This variable captures the spatial variation in trophic resource availability across the basin.

*Mixed Layer Depth* and *Sea Water Velocity* showed comparatively lower but still relevant importance, indicating the influence of physical oceanographic processes on prey aggregation and foraging opportunities. Cetaceans are often observed foraging along current boundaries where physical processes concentrate plankton and fish.

### 3.5. Species-Specific Variable Importance Patterns

While the community-level analysis revealed the overall ranking of environmental drivers, examination of species-specific variable importance patterns (Table 4) provides insights into niche differentiation and ecological segregation among the three species.

**Table 4.**
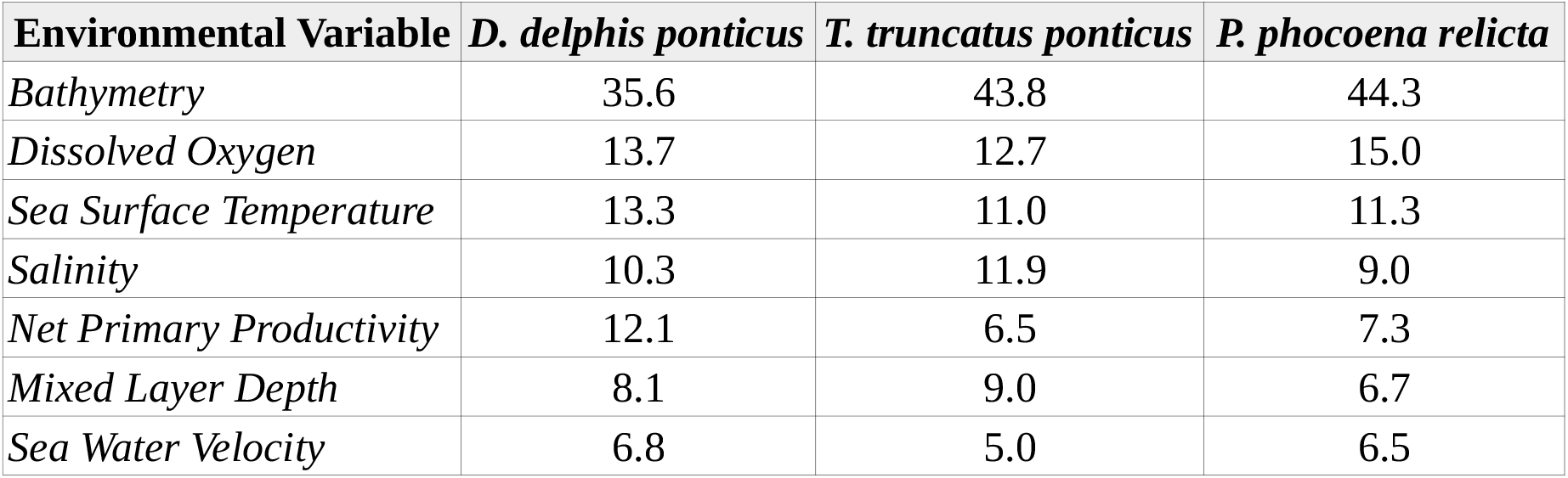
Species-specific variable importance rankings (% contribution)

The species-specific rankings reveal distinct environmental preferences consistent with known ecological traits (Sánchez-Cabanes et al., 2017):

- *D. delphis ponticus* (Common Dolphin). Bathymetry and Sea Surface Temperature were the primary drivers, reflecting its preference for deeper, warmer offshore waters. Primary Productivity also showed high importance, indicating reliance on productive foraging areas.
- *T. truncatus ponticus* (Bottlenose Dolphin). Bathymetry dominated the model, reflecting its strong association with shallower coastal and shelf waters. Salinity and Mixed Layer Depth showed elevated importance, consistent with its occurrence in estuarine and coastal areas influenced by freshwater inputs.
- *P. phocoena relicta* (Harbour Porpoise). Bathymetry and Dissolved Oxygen were the most important predictors, reflecting its preference for shallower, well-oxygenated shelf waters. Primary Productivity also ranked highly, indicating selection for productive foraging areas.

The distinct importance profiles support the hypothesis of niche segregation as a mechanism facilitating coexistence among the three sympatric cetacean species in the Black Sea. This pattern of differential environmental preferences reduces interspecific competition and allows the three species to partition available habitat resources.

## 4. Summary of Key Results

1. Individual ensemble species distribution models (ESDMs) for all three cetacean species achieved excellent predictive performance (AUC: 0.82–0.83; TSS: 0.65–0.67; prop.correct: 0.82–0.83).
2. The *Probability Ranking Rule (PRR)* method substantially improved stacked model performance (prediction.success = 0.459, sensitivity = 0.704, Jaccard = 0.465) compared to the uncorrected *pSSDM* (prediction.success = 0.36), demonstrating the value of overprediction correction in SSDM applications.
3. Species richness mapping identified the southwestern Black Sea shelf, Crimean coastal waters, the Kerch Strait region, and coastal areas nearby Georgia as high-richness areas of conservation priority.
4. Variable importance ranking revealed bathymetry as the most important community-level driver, followed by dissolved oxygen, sea surface temperature, and salinity.
5. Species-specific importance patterns confirmed ecological niche segregation, with common dolphins favouring deeper offshore waters, while bottlenose dolphins and harbour porpoises are more strongly associated with shallower shelf environments.

